# ISCA2 inhibition decreases HIF and induces ferroptosis in clear cell renal carcinoma

**DOI:** 10.1101/2022.06.01.494206

**Authors:** Yangsook Song Green, Maria C. Ferreira dos Santos, Daniel Fuja, Ethan Reichert, Alexandre R. Campos, Sophie J. Cowman, Jessica Kohan, Sheryl R. Tripp, Elizabeth A. Leibold, Deepika Sirohi, Neeraj Agarwal, Xiaohui Liu, Mei Yee Koh

**Author notes:** Corresponding author Corresponding address: Mei Yee Koh, Department of Pharmacology and Toxicology, University of Utah, 30S 2000E, Salt Lake City, UT 84112. Contributed equally (Yangsook Song Green, Maria C. Ferreira dos Santos, Daniel Fuja).

## Abstract

Clear cell renal cell carcinoma (ccRCC), the most common form of kidney cancer, is typically initiated by inactivation of the von Hippel Lindau (*VHL)* gene, which results in the constitutive activation of the hypoxia inducible factors, HIF-1α and HIF-2α. Using a high throughput screen, we identify novel compounds that decrease HIF-1/2α levels and induce ferroptosis by targeting Iron Sulfur Cluster Assembly 2 (ISCA2), a component of the late mitochondrial Iron Sulfur Cluster (L-ISC) assembly complex. ISCA2 inhibition either pharmacologically or using siRNA decreases HIF-2α protein levels by blocking iron-responsive element (IRE)-dependent translation, and at higher concentrations, also decreases HIF-1α translation through unknown mechanisms. ISCA2 inhibition also triggers the iron starvation response, resulting in iron/metals overload and death via ferroptosis. ISCA2 levels are decreased in ccRCC compared to normal kidney, and decreased ISCA2 levels are associated with pVHL loss and with sensitivity to ISCA2 inhibition. Strikingly, pharmacological inhibition of ISCA2 using an orally available ISCA2 inhibitor significantly reduced ccRCC xenograft growth *in vivo*, decreased HIF-α levels and increased lipid peroxidation, suggesting increased ferroptosis *in vivo*. Thus, the targeting of ISCA2 may be a promising therapeutic strategy to inhibit HIF-1/2α and to induce ferroptosis in pVHL deficient cells.

## Introduction

Clear cell renal cell carcinoma (ccRCC) is highly refractory to standard chemotherapy and radiation and is one of the most common and aggressive subtypes of kidney cancer. Patients with advanced or metastatic tumors (30% of patients) have a 5-year survival rate of just 13% [1]. The etiology of ccRCC is uniquely linked to loss of the von Hippel Lindau (*VHL*) tumor suppressor gene, resulting in the pseudo-hypoxic activation of the hypoxia-inducible factors, (HIF)-1α and HIF-2α [2, 3] [4]. The VHL protein (pVHL) is the substrate recognition component of the E3 ligase complex that targets HIF-1α and HIF-2α subunits for proteasomal degradation under aerobic conditions [5]. Under hypoxic conditions, or in the presence of pVHL loss-of-function mutations, HIF-1α and HIF-2α are stabilized, and enter the nucleus where they heterodimerize with HIF-1β, forming the HIF-1 or HIF-2 transcriptional complexes, respectively. The HIF heterodimers bind to conserved hypoxia response elements (HREs) within regulatory regions of target genes to activate transcription of hundreds of genes critical for the adaptation to hypoxia, and for tumor progression, such as those promoting aerobic glycolysis, angiogenesis, and metastasis [6, 7]. In addition to canonical HRE-mediated transcription requiring hetero-dimerization with HIF-1β, the HIF-1α and HIF-2α subunits differentially modulate cellular signaling pathways through interaction with proteins that do not contain PAS domains, including the tumor suppressor protein p53, the c-MYC proto-oncogene, β-catenin and the Notch intracellular domain [8-13].

HIF-1 versus HIF-2-specific activation is determined, at least in part, by factors within the tumor microenvironment such as intensity and duration of hypoxia, as well as cell-type-specific pathway activation within both tumor cells and those of the complex tumor microenvironment [14-16]. Consequently, although the HIFs share many transcriptional targets, they also regulate non-overlapping genes: For example, anerobic glycolysis is predominantly HIF-1 controlled, whereas erythropoietin *(EPO)* synthesis and iron uptake in the gut have emerged as primarily HIF-2-regulated processes [17-21]. Indeed, HIF-2α plays a non-redundant role in iron regulation and is additionally regulated at the translational level by an RNA stem-loop element known as an iron-responsive element (IRE), within the 5’ untranslated region (UTR) of the HIF-2α transcript [22]. Under conditions of iron depletion, iron regulatory proteins (IRP1 and IRP2) bind to IREs within 5′ or 3’ UTRs of transcripts resulting in translational repression and transcript stabilization respectively. The IRPs coordinate the cellular response to iron starvation by decreasing iron storage and increasing iron uptake such as through downregulation of the central iron storage molecule, ferritin (FTH1; 5’IRE) and upregulation of the major mediator of cellular iron uptake, transferrin receptor (TFRC; 3’IRE) respectively. In this regard, binding of IRP1 to the 5’IRE of HIF-2α mediates HIF-2α translational repression during conditions of iron (or Fe-S cluster) insufficiency to limit erythropoiesis [22].

Despite significant homology in the sequence, structure, and regulation of HIF-1α and HIF-2α, HIF-2α has been proposed to play a dominant driving role over HIF-1α in ccRCC. [23, 24]. The HIF-2 bias observed in ccRCC may be due to the increased potency of HIF-2α compared to HIF-1α in promoting pro-tumorigenic factors such as Cyclin D1, TGF-α and VEGFA ^[23, 25]^. HIF-2α has also been implicated in ccRCC metastasis through activation of CXCR4, and in driving the transcription of stem cell factors *OCT-3/4* and *SOX2*, which promote an undifferentiated, more aggressive phenotype [26-31]. Indeed, the first-in-class HIF-2α selective inhibitor, belzutifan, which blocks the heterodimerization of HIF-2α with HIF-1β, shows promising single-agent activity in heavily pre-treated patients with advanced ccRCC, and was recently approved for the treatment of cancers associated with VHL disease including ccRCC [32, 33]. Although these studies support a tumor-promoting role for HIF-2α in ccRCC, recent data suggest that HIF-1α may also contribute to ccRCC progression in the context of the complex tumor microenvironment [15, 34, 35]. This suggests that the targeting of HIF-1α, in addition to HIF-2α may provide additional therapeutic benefit in ccRCC.

In this study, we describe the results of a high-throughput screen for selective inhibitors for HIF-2α that reveal a series of small molecules that decrease HIF-2α protein, and at higher concentrations, also HIF-1α. These compounds mediate their effects by inhibiting the function of iron sulfur cluster assembly 2 (ISCA2) which blocks IRE-dependent HIF-2α translation and inhibits HIF-1α translation through unknown mechanisms. Significantly, we show that these ISCA2 inhibitors trigger the iron starvation response resulting in the accumulation of iron (and other transition metals) that induce death via ferroptosis both *in vitro* and *in vivo*. Ferroptosis is a form of necrotic cell death associated with iron-dependent oxidation of phospholipid membranes, which causes defects within the plasma and mitochondrial membranes resulting in cell death [36, 37]. The induction of ferroptosis is a promising therapeutic strategy for cancer, particularly those with clear cell morphology such as ccRCC [38]. A number of potent ferroptosis inducers that trigger ferroptosis *in vitro* such as by depleting intracellular glutathione (e.g. erastin) or inhibiting GPX4 (e.g. RSL3) have been described but these are unsuitable as clinical candidates, at least in part, due to their unfavorable drug-like properties that make them unsuitable for oral administration [39, 40].

Here, we describe a novel role of ISCA2 in HIF and iron regulation and highlight its value as a potential therapeutic target in ccRCC for the dual inhibition of HIF-α and the induction of ferroptosis.

## Results

### A high-throughput screen for HIF-2α inhibitors identify compounds that decrease HIF-2α translation

To identify inhibitors of HIF-2α, we performed a high throughput screening campaign of the 360,000 compound NIH Molecular Libraries Probe Production Centers Network (MLPCN) using a hypoxia-responsive element (HRE)-luciferase based screen in 786-0 ccRCC cells (express HIF-2α exclusively), followed by a counter-screen using deferoxamine-treated MIA-PaCa2 pancreatic cancer cells (express HIF-1α exclusively). Screen flow chart and HIF-α dependency of HRE-luciferase activity in 786-0 HRE and MIAPaCA-2 HRE cells are shown in **Fig. 1A-C** respectively. In the initial screen, 2626 compounds were identified as primary actives (Z’ average = 0.84, Signal/background ratio = 76.8) in 786-0 HRE cells. Promiscuous compounds were filtered, reducing the number of actives to 1927. Of these, 982 hits were reconfirmed using fresh powder. These compounds were counter-screened in the MIAPaCA-2 HRE cells, identifying 209 compounds producing ≤50% activity. Retesting with fresh powder and eliminating non-actives or non-selective inhibitors of luciferase, followed by structure activity relationship (SAR) studies identified compound #1 and its more potent derivative, compound #25 that decreased HIF-2α activity and protein in a variety of cell lines (**Fig. 1D-G, Supplemental data S1A**). These compounds also decreased HIF-1α levels at higher concentrations or treatment durations (**Fig. 1F-G**). Treatment with the proteasomal inhibitor, MG132 did not prevent the decrease in HIF-1/2α mediated by #1 or #25 (**Fig. 1G**), suggesting that these compounds do not promote proteasomal degradation of HIF-1/2α. Furthermore, although treatment of cells with #1 or #25 resulted in a dose-dependent decrease in the transcription of HIF-2α target genes *VEGFA* and *OCT-3/4*, neither compound significantly affected levels of HIF-2^α^ (*EPAS1*) transcript (**Fig. 1H, S1B**), indicating that these compounds do not inhibit HIF-2α transcription. Indeed, we found that #1 and #25 inhibited the production of luciferase driven by the HIF-2α IRE (IRE-Luc) (**Fig. 1I, S1C**) suggesting that these compounds block IRE-dependent translation of HIF-2α [22]. The decrease in HIF-2α translation was confirmed by decreased puromycin incorporation into immunoprecipitated HIF-2α in hypoxic ACHN cells treated with #1 using the SUnSET assay (**S1D**) [41]. In addition to decreasing HIF1/2α levels, we also noted dose dependent increases in IRP2 and TFRC and decreased FTH after treatment with #1 or #25, which together indicate the triggering of the iron starvation response, which would be consistent with the IRP1/IRE mediated translational inhibition of HIF-2α that we have observed (**Fig. 1E-F**). Thus, using a high-throughput screening campaign, we have identified a series of small molecules that decrease HIF-2α by inhibiting IRE-dependent translation, and that also inhibit HIF-1α at higher concentrations/treatment durations.

**Figure 1:**
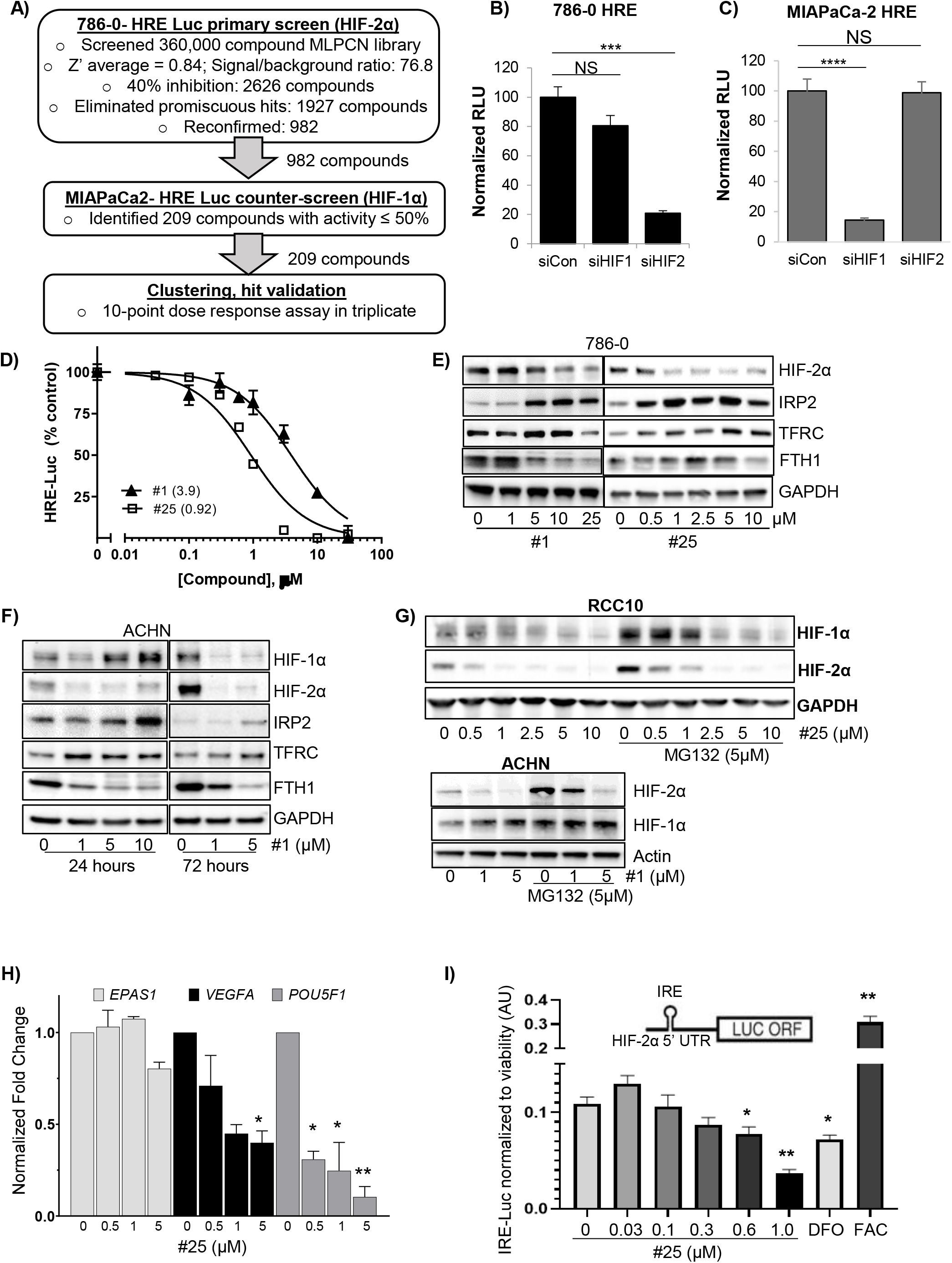
Identification and characterization of novel HIF-2α selective inhibitors. A) Screening flow-chart for the identification of selective small molecule inhibitors of HIF-2α. B-C) Validation of HIF-1α or HIF-2α dependency of a Hypoxia Responsive Element (HRE)-driven luciferase reporter construct stably expressed in (B) 786-0 or (C) MIAPaCa-2 cells. D) 786-0 HRE-Luciferase dose-response assays of indicated compounds (24-hour treatment) with IC_50_ values (μM) indicated in brackets. E) Western blots showing effects of 24-hour treatment with indicated concentrations of compounds (μM) on levels of HIF-2α and other proteins in 786-0 cells in normoxia. F) Effects of 24- or 72-hours’ treatment with #1 in ACHN cells (exposed to hypoxia for the final 24 hours). G) Effects of treatment of, (top) RCC10 cells with #25 ± proteasomal inhibitor, MG132 (24 hours, 20%O_2_), or of (bottom) hypoxic ACHN cells #1 ± MG132 (24 hours, 1%O_2_) on HIF-1α and HIF-2α. H) Quantitative RT-PCR showing effects of #25 on the transcription of *HIF2A* (*EPAS1*) and HIF-2 target genes *VEGFA* and *OCT-3/4* (*POU5F1*). I) Effect of #25 treatment on luciferase activity driven by a HIF-2α Iron-Responsive Element (IRE)-luciferase reporter construct stably expressed in 786-0 cells. The iron chelator, deferoxamine (DFO; 50μM), and iron donor, ferric acetylcysteine (FAC; 25µM) were used to confirm responsiveness of the reporter to iron perturbation. All data shown are representative of at least 2 independent experiments and show the mean ± SEM. * p <0.05; ** p < 0.01.

### HIF-α inhibitors target ISCA2 and trigger the iron starvation response

To identify the molecular target of #1 and #25, we used the drug affinity responsive target stability (DARTS) assay, which is based on the principle that the binding of a small molecule to its target renders the latter protease resistant [42]. We used 2D-LC-MS/MS to identify peptides differentially degraded by pronase (a mixture of proteases) in the absence or presence of #1 (100µM) using hypoxic ACHN cell lysates (**Fig. 2A**). From >2000 unique proteins detected, we confirmed ISCA2 as the most likely molecular target for #1 via both 2D-LCMS/MS and western blotting (**Fig. 2B**). Indeed, pre-treatment with #1 prior to pronase incubation increased the amount of ISCA2 detected by 2D-LC-MS/MS by almost 5-fold, and this increase in ISCA2 levels were detectable via western blotting (**Fig. 2B**), suggesting that #1 protects ISCA2 from pronase-mediated degradation. ISCA2 was further validated as the likely molecular target of these compounds using thermal shift studies using recombinant ISCA2 wherein treatment with either #1 or #25 induced significant increases in the melting temperature of ISCA2 compared to DMSO alone (**Fig. 2B-C**). siRNA knockdown of ISCA2 dramatically reduced both HIF-1α and HIF-2α protein levels, confirming ISCA2 as the likely molecular target of these compounds (**Fig. 1D**). However, unlike treatment with #1 and #25 which induced IRP2 and TFRC, ISCA2 siRNA markedly reduced the levels of these proteins (**Fig. 1D**), which may be due to the extended duration of ISCA2 siRNA transfection required to reduce ISCA2 protein levels (6 days), compared to the relatively short period of drug treatment (24 or 72 hours). Regardless of these differences, we identified a significant increase in iron and other transition metals when cells were treatment with compound #1 or #25 or transfected with ISCA2 siRNA (**Fig. 2E-G, S2A-C**). Thus, the data suggest that these compounds decrease HIF-1/2α levels and induce iron and metals accumulation by targeting ISCA2.

**Figure 2:**
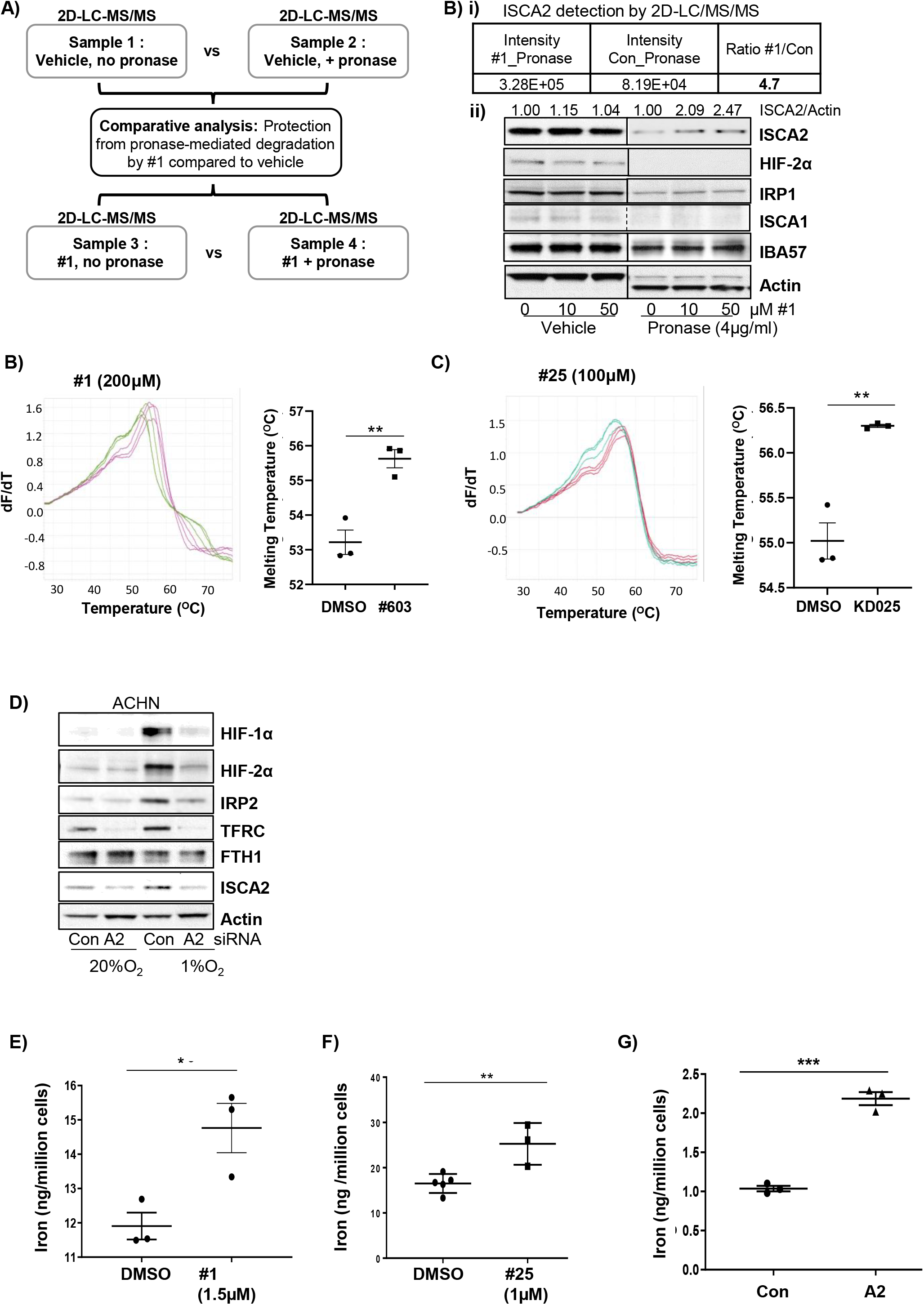
ISCA2 is the molecular target of HIF-α inhibitors. A) Schematic showing sample preparation workflow for DARTS assay. Hypoxic ACHN lysates were split into 4 samples and subjected to indicated treatments. B) i) Mass spectrometry data from DARTS assay indicating intensity of ISCA2 fragments in cell lysates treated with Pronase together with #1 or DMSO (Con); ii) Western blot showing effects of control or Pronase treatment in hypoxic ACHN cell lysates treated with indicated concentrations of #1. Densitometric analysis of ISCA2 band intensities normalized to actin is shown above blots. B-C) Thermal shift assays showing effects of incubation of recombinant ISCA2 (4μg) with (B) DMSO (green) or 200μM #1 (magenta), or (C) DMSO (green) or 100µM #25 (red). Quantitation of ISCA2 melting temperatures are shown inset (mean ± SEM). D) Effects of transfections with non-targeting (Con) or ISCA2 (A2) siRNA on indicated proteins in hypoxic ACHN cells. Knockdown was performed using two consecutive siRNA transfections over 72 hours each. E-G) Iron content of 786-0 cells treated with (E) #1, (F) #25, both for 24 hours; or (G) transfected with siCon or siISCA2 (6 days) as determined using ICP-MS.

### ISCA2 mediates its effects on HIF regulation and iron sensing in ways that are independent of its role in the mitochondrial iron sulfur cluster assembly complex

ISCA2, together with ISCA1 and IBA57 (Iron-Sulfur Cluster Assembly Factor for Biotin Synthase-And Aconitase-Like Mitochondrial proteins) is a component of the late mitochondrial Iron-Sulfur Cluster Assembly (L-ISC) complex, which regulates the maturation of mitochondrial iron-sulfur [4Fe-4S] proteins, which includes the respiratory chain complexes I and II, mitochondrial aconitase and lipoic acid synthase [43-46]. To determine whether knockdown of the other components of the L-ISC caused similar effects to ISCA2 on HIF regulation and iron sensing, we compared the effects of knockdown of each of the components of the L-ISC. Intriguingly, we found that only knockdown of ISCA2 decreased levels of HIF-1α and HIF-2α, and decreased levels of IRP1, IRP2 and TFRC in ACHN cells 6 days post-siRNA transfection (**Fig. 3A**). We also observed decreased levels of IBA57 with ISCA2 siRNA, suggesting that the stability of IBA57 may be regulated by ISCA2. The effects on ISCA2 knockdown on the HIFs were recapitulated using a different siRNA for ISCA2, showing that HIF-1α and HIF-2α levels were decreased as soon as 2 days post-transfection, when the decrease in ISCA2 protein (and in IRP1, IRP2 and TFRC) was less apparent (**Fig. 3B**). We also observed a rapid decrease in FTH1 that was not observed after 6 days’ siRNA transfection, which resembles that seen when cells were treated with #1 and #25 (**Fig. 3B, 1E-F**). Similar effects in decreasing HIF-1/2α, IRP1, IRP2 and TFRC were obtained in RCC4 cells which are pVHL deficient, suggesting that ISCA2 plays a unique role in HIF regulation and iron sensing independently of its role in the L-ISC and independently of pVHL in multiple cell lines (**Fig. 3C**). As seen with #1 and #25 (**Fig. 1G**), co-treatment of siRNA-transfected ACHN cells with MG132 did not prevent the loss of HIF-α associated with ISCA2 knockdown (**Fig. 3D**). Intriguingly, unlike that seen with #1 and #25, siRNA-mediated knockdown of ISCA1, ISCA2 and IBA57 significantly decreased *EPAS1* transcription but did not significantly affect *HIF1A* transcription (**Fig. 3E**). Since both ISCA1 and IBA57 knockdown decreased *EPAS1* transcript without affecting HIF-2α protein levels in ACHN cells, it is unlikely that the HIF-2α decrease mediated by ISCA2 siRNA is primarily due to decreased transcription. However, the decreased *EPAS1* transcription may be due to mitochondrial retrograde signaling in which dysfunctional mitochondria communicate with nuclear components, typically modulating transcription in response to mitochondrial stresses [47]. Using the SunSET assay, we found that ISCA2 knockdown abrogated the incorporation of puromycin into immunoprecipitated HIF2α, consistent with IRE-mediated inhibition of translation (**Fig. 3F**). Intriguingly, we observed similar effects on of ISCA2 knockdown on HIF-1α translation, suggesting that ISCA2 may regulate HIF-1α translation through an unknown mechanism (**Fig. 3F**). A robust readout of mitochondrial [4Fe-4S] function is the mitochondrial [4Fe-4S] dependent lipoylation of dihydrolipoamide S-acetyltransferase (DLAT), and dihydrolipoamide S-succinyltransferase (DLST) [44, 48]. Knockdown of either ISCA1 or ISCA2 markedly decreased the lipoylation of DLAT and DLST in HeLa and 786-0 cells, whereas knockdown of IBA57 had minimal effects on lipoylation, confirming that both ISCA1 and ISCA2 play essential roles mitochondrial [4Fe-4S] protein assembly (**S3A**). Treatment of HeLa or 786-0 cells with #1 or #25 respectively similarly inhibited the lipoylation of DLAT and DLST suggesting that these compounds also inhibit mitochondria [4Fe-4S] assembly (**Fig. 3G**) consistent with their functions as inhibitors of ISCA2. However, when we investigated the impact of the ISC components on iron/metals content, we found that only ISCA2 knockdown induced marked elevation of total cellular iron and of other transition metals in ACHN and 786-0 cells (**Fig. 3H, S3B**). Taken together, our data suggest that ISCA2 knockdown or pharmacological inhibition decreases the translation of HIF-1α and HIF-2α and promotes metals accumulation through a mechanism that is independent of ISCA2’s role in the L-ISC.

**Figure 3:**
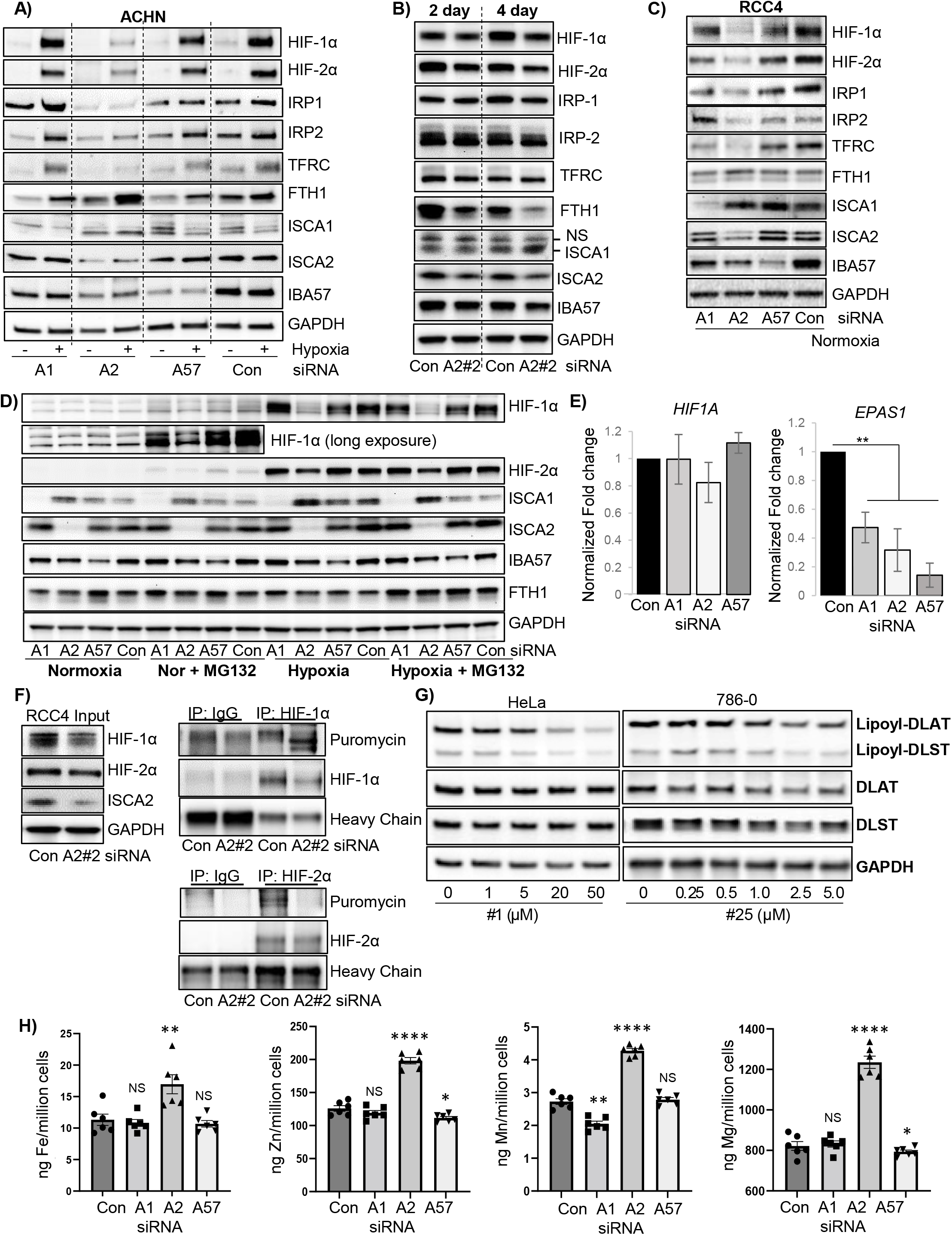
ISCA2 depletion decreases HIF-1/2α levels and promotes iron and metals accumulation. A Western blots showing effect of 6-day siRNA knockdown of ISCA1 (A1), ISCA2 (A2#1), IBA57 (A57) or non-targeting siRNA (Con) in ACHN cells in normoxia or after 24 hours at 1% O_2_ (hypoxia). B) Effect of 2- and 4-day knockdown of ISCA2 using a second siRNA construct (A2 #2) in ACHN cells in hypoxia. C) Effect of 6-day knockdown with indicated siRNAs in RCC4 cells. D) Effect of MG132 on ACHN cells transfected indicated siRNAs for 6 days in normoxia or hypoxia (5μM MG132 in normoxia and 1μM MG132 in hypoxia, MG and hypoxia exposure performed for the last 6 hours). E) Effect of transfection of indicated siRNAs on the transcription of *HIF1A* and *EPAS1* normalized to *B2M* in hypoxic ACHN cells. F) Effect of ISCA2 siRNA on HIF-1/2α translation in RCC4 cells. Cells were transfected with ISCA2 siRNA for 3 days, then labelled with puromycin. Puromycin incorporation was determined by immunoprecipitation of HIF-1/2α followed by western blot using an antibody against puromycin. G) Effect of #1 or #25 treatment (24 hours) on lipoylation of DLAT and DLST in HeLa and 786-0 cells respectively. H) Effect of 5-day siRNA knockdowns on metals accumulation in ACHN cells detected using ICP-MS. Approximately equal cell numbers were submitted for analysis. All data are representative or averages of at least two independent experiments shown as mean ± SEM.

### ISCA2 inhibition promotes cell death via ferroptosis

We noted that when performing the siRNA knockdown of ISCA1, ISCA2 and IBA57 that ISCA2 knockdown consistently decreased protein content to a greater extent than that observed with ISCA1 or IBA57 knockdown in a variety of cell lines (**Fig. 4A**). We also noted a consistent difference in media color indicating a higher pH in cells transfected with ISCA2 compared to that of Con, ISCA1 or IBA57 siRNA transfected cells, which may reflect the strikingly lower cell density (and hence reduced metabolic activity) associated with ISCA2 knockdown (**Fig. 4B, S4**). In this regard, only ISCA1 knockdown promoted acidification of cell culture media, which likely reflects a defect in respiratory function (**Fig. 4B**) as previously reported [44]. Indeed, the profound reduction in the number of viable cells obtained with ISCA2 knockdown was confirmed using manual counting of trypan blue-excluding viable cells using two separate siRNAs (**Fig. 4C**). Cells transfected with ISCA2 siRNA also exhibited necrotic morphology such as cell enlargement and plasma membrane rupture, commonly observed in ferroptotic cells (**S5**) [49]. ISCA2 knockdown also significantly decreased the number of viable 786-0 cells compared to those transfected with Con siRNA, ISCA1 or IBA57 siRNA using the resazurin cell viability (**Fig. 4D**). Although no markers exist that can distinguish ferroptosis from other forms of programmed cell death, we used a kit that enables real-time monitoring of changes associated with cell death (Promega JA1011). We show that siRNA knockdown of ISCA2 resulted in induction of phosphatidyl serine (PS) on the cell surface, typically associated with early apoptosis, 28 hours after siRNA transfection (**Fig. 4E**) followed by loss of membrane permeability, typically associated with secondary necrosis, 72 hours after siRNA transfection (**Fig. 4F**). Treatment of 786-0 cells with #1 or #25 similarly decreased cell viability, and this could be attenuated by co-treatment with the iron chelator, DFO, the ferroptosis inhibitor, liproxstatin, or the free radical scavenger N-acetyl cysteine (NAC), but not with the caspase inhibitor ZVAD-FMK, suggesting that ISCA2 inhibition promotes death via ferroptosis associated with iron/metals overload (**Fig. 4G-H**). Furthermore, #25 initially induced cell surface exposure of PS (**Fig. 4I**, left Y-axis), followed by loss of membrane permeability (**Fig. 4I**, right Y-axis), which was markedly attenuated by co-treatment with liproxstatin (**S6A**), in a time course effect that closely resembled that of the GPX4 inhibitor RSL3, suggesting that the ISCA2 inhibitors induce ferroptosis with characteristics and kinetics resembling that of RSL3. We also noted that treatment with higher concentrations of #25 markedly decreased levels of GPX4 (**S6B**), which may sensitize cells to ferroptosis. We found that #25 treatment increased levels of the oxidized form of C11-BODIY (581/591), a fluorescent probe for indexing lipid peroxidation, confirming that #25 induces the lipid peroxidation associated with ferroptosis (**Fig. 5A**) [50]. Additionally, treatment with #25 but not with PT2385 (an analog of belzutifan) increased levels of malondialdehyde (MDA)-thiobarbituric (TBA) adducts, another indicator of lipid peroxidation, to a level comparable to the GPX4 inhibitor, RSL3, further confirming the ISCA2 inhibitors as inducers of lipid peroxidation and downstream ferroptosis (**Fig. 5B**) [51]. Taken together, the data suggests that ISCA2 inhibition promotes cell death via ferroptosis independently of its role in the ISC complex.

**Figure 4:**
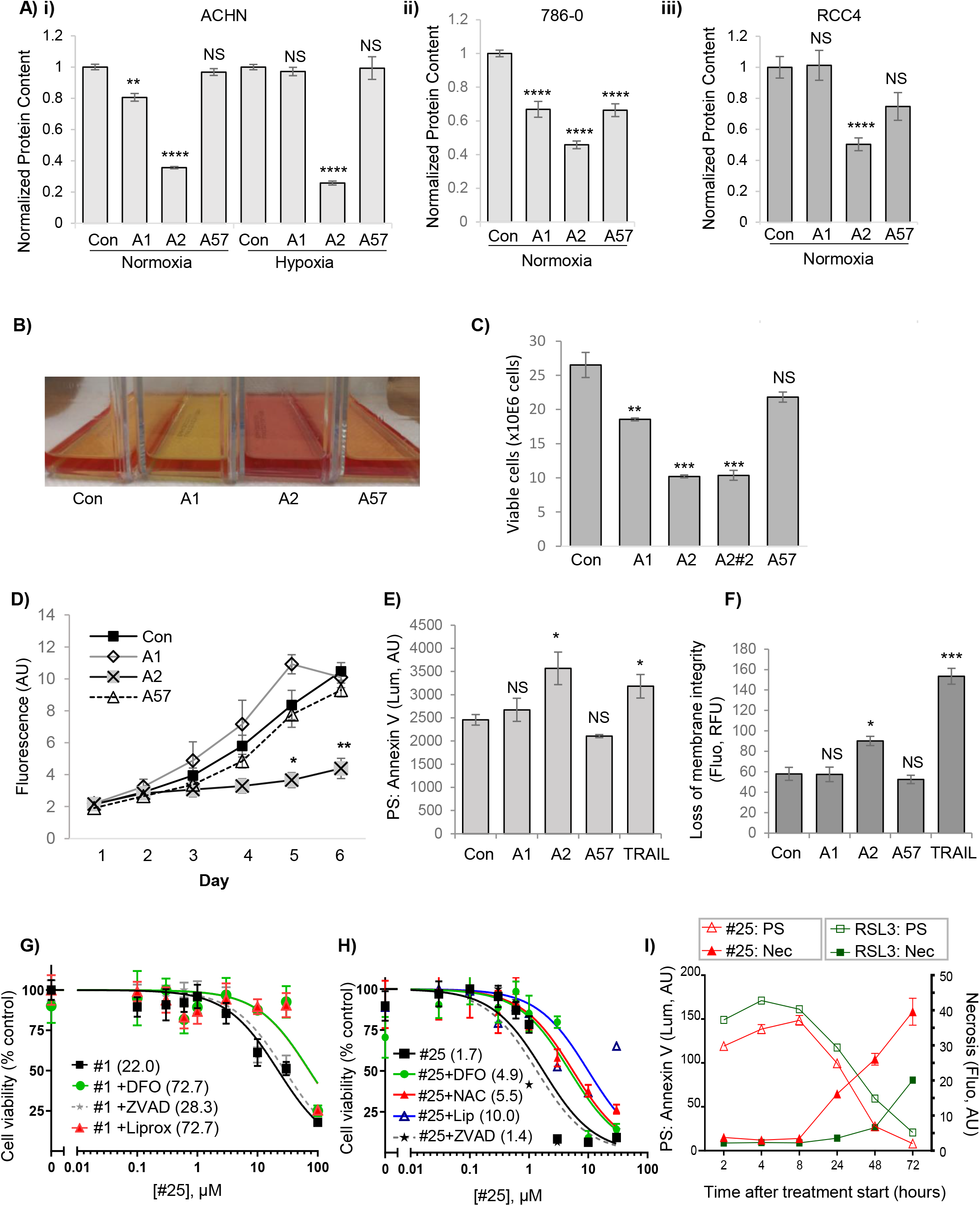
SiRNA knockdown or pharmacological inhibition of ISCA2 induces ferroptosis. A) Effect of indicated siRNAs on total protein content in indicated cell lines (6 days post-transfection). B) Media appearance of ACHN cells transfected with indicated siRNAs for 5 days. C) Viable cell number determined by trypan blue exclusion in ACHN cells 5 days after transfection with indicated siRNAs. D) Cell viability assay of 786-0 cells transfected with indicated siRNAs. Cells were seeded into 96-well plates 6 days after siRNA transfection, and cell viability was measured using resazurin daily for 6 days. E) Effect of indicated siRNA on PS exposure in 786-0 cells 28 hours after transfection. TRAIL (400ng/ml, 4hrs) is a positive control. F) Effect of siRNA transfection in causing loss of membrane permeability in 786-0 cells 72hrs after transfection using TRAIL (400ng/ml, 72hrs) as a positive control. G-H) Cell viability assays (resazurin) of 786-0 cells treated with (G) #1 ± DFO (100μM), liproxstatin (1μM) or ZVAD-FMK (20μM; Or (H) #25 ± DFO (100μM), liproxstatin (1μM), NAC (1mM) or ZVAD-FMK (20μM). Treatments were performed for 24 hours. Average IC_50_ values (µM) are shown in brackets. I) Time course of PS: annexin V binding (left Y-axis) and loss of membrane permeability (i.e. necrosis - right Y-axis) in RCC10 cells after #25 (3μM) or RSL3 (0.5µM) treatment. Data shown are the mean ± SD.

**Figure 5:**
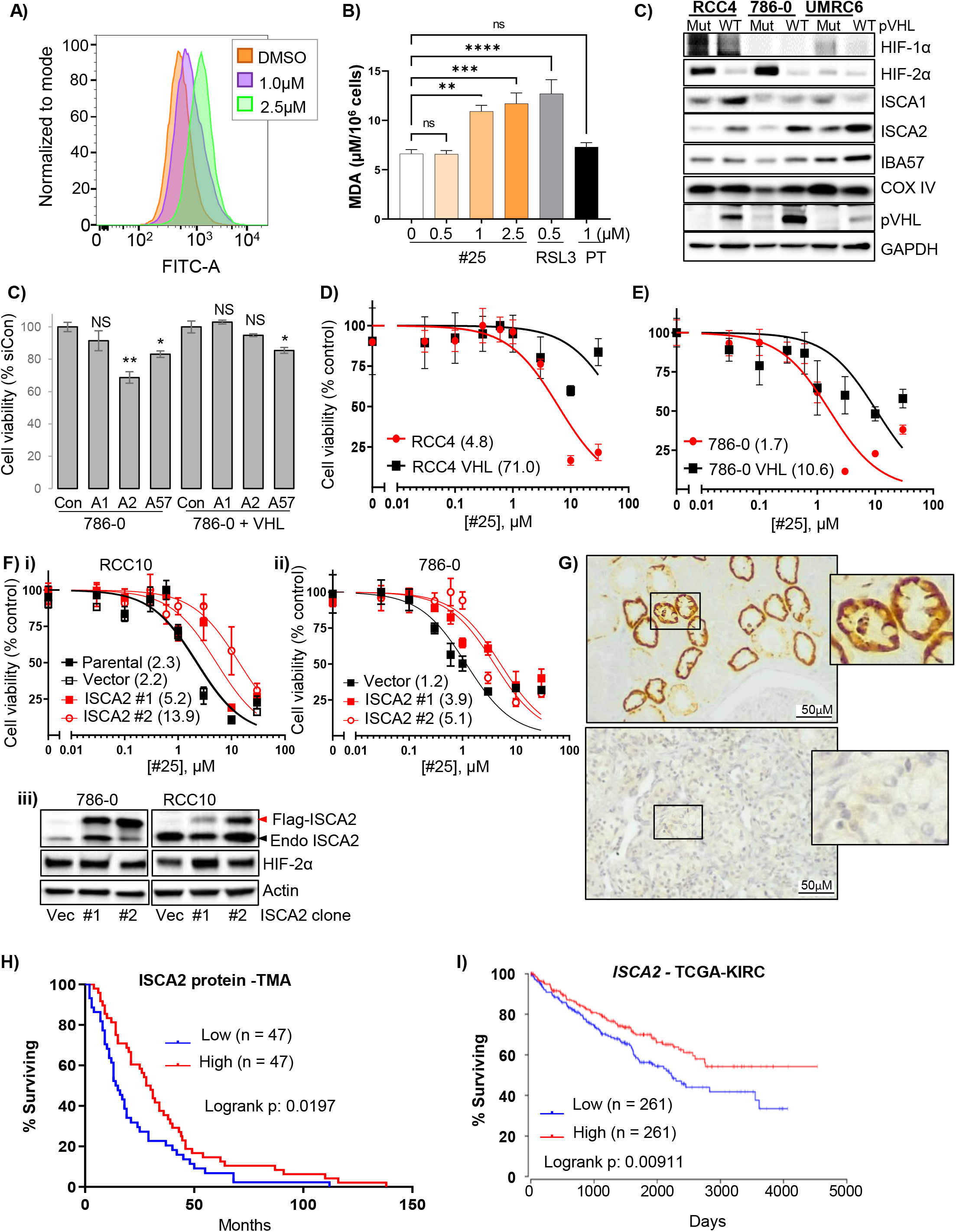
ISCA2 inhibition promotes lipid peroxidation and ferroptosis and low ISCA2 levels are associated with poor prognosis. A) Impact of #25 treatment (48 hours) on BODIPY 581/591 fluorescence detected by flow cytometry in 786-0 cells. B) Impact of 48 hours’ treatment with #25, PT2385 (PT) or 6 hours’ treatment with RSL3 on malondialdehyde (MDA), an indicator of lipid peroxidation in RCC10 cells. C) ISCA2 levels in pVHL mutant ccRCC cells ± stable pVHL reconstitution. D) Resazurin cell viability assay of 786-0 parental or 786-0 cells with pVHL reconstitution transfected with indicated siRNAs (24 hours post-transfection). D-E) Representative plot showing viability (resazurin) of RCC4 and RCC4 + VHL cells (D), and 786-0 and 786-0 +VHL cells (E), after treatment with #25 for 24 hours. IC_50_ values (μM) are shown in brackets. F) Representative plots showing viability of RCC10 (i), or 786-0 (ii) cells stably expressing empty vector (Vec) or ISCA2 (2 clones shown) treated with #25 for 24 hours. IC_50_ values (μM) are shown in brackets. Western blots showing endogenous (Endo) and overexpressed FLAG-ISCA2 levels are shown in (iii). G) Immunohistochemistry showing representative expression of ISCA2 in normal kidney (top) and in ccRCC (bottom). H, I) Kaplan-Meier curves showing associations of (H) ISCA2 protein, or (I) *ISCA2* transcript levels above (high) or below (low) the median with overall survival using a tumor microarray (TMA; 94 cases) or data from TCGA-KIRC (522 cases) respectively

### ISCA2 levels are increased with pVHL reconstitution is associated with better patient prognosis

To investigate the relationship between ISCA2 and pVHL, we examined the effect of stable pVHL reconstitution in a panel of pVHL-deficient ccRCC cells. Strikingly, pVHL reconstitution resulted in marked upregulation of ISCA2 protein, whereas levels of ISCA1, IBA57 and COX-IV did not show major changes (**Fig. 5C**), suggesting that the ISCA2 increase may not simply be a product of increased mitochondrial mass that is associated with pVHL re-expression [52-56]. Intriguingly, ISCA2 knockdown resulted in a significant decrease in cell viability in 786-0 but not in 786-0 VHL cells relative to siCon (**Fig 5C**). pVHL re-expression was also associated with a marked increase in resistance to #25, as cells with pVHL reconstitution showed approximately 6- and 15-fold increases in #25 viability IC_50_s in RCC4 and 786-0 VHL cells compared to their parental pVHL deficient cells, respectively (**Fig. 5D-E**) with similar results seen with #1 (**S6C**). Similarly, stable ISCA2 overexpression was also associated with marked increases in resistance of RCC10 and 786-0 cells to ISCA2 inhibition by #25 (**Fig. F**), supporting the notion that cells with low ISCA2 are more sensitive to ISCA2 inhibition than those with high ISCA2. To investigate the expression of ISCA2 in human tissue, we stained sections of ccRCC and uninvolved normal kidney for ISCA2. Analysis by board certified genitourinary pathologist (DS) revealed that ISCA2 staining in normal kidney exhibited a primarily diffuse cytosolic localization (which cannot be used to distinguish mitochondrial from cytosolic localization), with most intense staining within the epithelial cells of the proximal convoluted tubule (PCT, **Fig. 5G**). By contrast, ISCA2 staining was markedly lower in ccRCC tissue compared to uninvolved kidney possibly due to oncogenic transformation of PCT and resultant loss of structural integrity of organelles in which ISCA2 is found (**Fig. 5G)**. To determine the relationship between ISCA2 levels and patient outcome, we investigated the association of ISCA2 protein levels and overall survival of in a cohort of 94 patients treated with anti-angiogenic tyrosine kinase inhibitors. Intriguingly, patients with ISCA2 levels above the median had significantly increased overall survival (hazard ratio high vs low ISCA2: 0.68; p = 0.0197; **Fig. 5H**) compared to those with ISCA2 levels below the median. We observed a similar association of high *ISCA2* transcript levels with better overall survival in TCGA-KIRC (**Fig. 5I**), which, taken together with our TMA data suggest that high ISCA2 levels are association with better outcome. Thus, the data suggest that low ISCA2 levels in ccRCC are associated with pVHL loss and with poorer prognosis, and that cells lacking functional pVHL are more sensitive to ISCA2 inhibition, possibly due to decreased levels of ISCA2. Taken together, the data suggest that the targeting of ISCA2 could be a strategy to inhibit HIF-α and induce ferroptosis particularly in pVHL deficient, more aggressive ccRCC cells.

### ISCA2 inhibition significantly decreases tumor growth in vivo

To investigate the feasibility of ISCA2 inhibition as a therapeutic strategy for ccRCC, we used compound #25 as a probe compound, based on its acceptable drug-like properties (bioavailability F = 27.3%; half-life T_1/2_: 3.61 hours, Cmax: 1913ng/ml, AUC_0-inf_:4161ng for 10mg/kg dose, not shown). Oral administration of #25 once daily by oral administration resulted in a significant dose-dependent reduction of 786-0 ccRCC tumor growth *in vivo* (**Fig. 6A**). Daily administration of #25 was well tolerated with no significant weight loss throughout the study (**Fig. 6B**). Treatment with 60mg/kg #25 in RENCA syngeneic tumor xenografts also significantly inhibited tumor growth (**Fig. 6C**). This was accompanied by significantly decreased HIF-1α (HIF-2α levels were undetectable in these tumors) and in significantly elevated MDA (**Fig. 6D-E**). Thus, ISCA2 inhibition decreases HIF-α levels and induces the lipid peroxidation associated with ferroptosis both *in vitro* and *in vivo* and is a promising treatment strategy for ccRCC.

**Figure 6:**
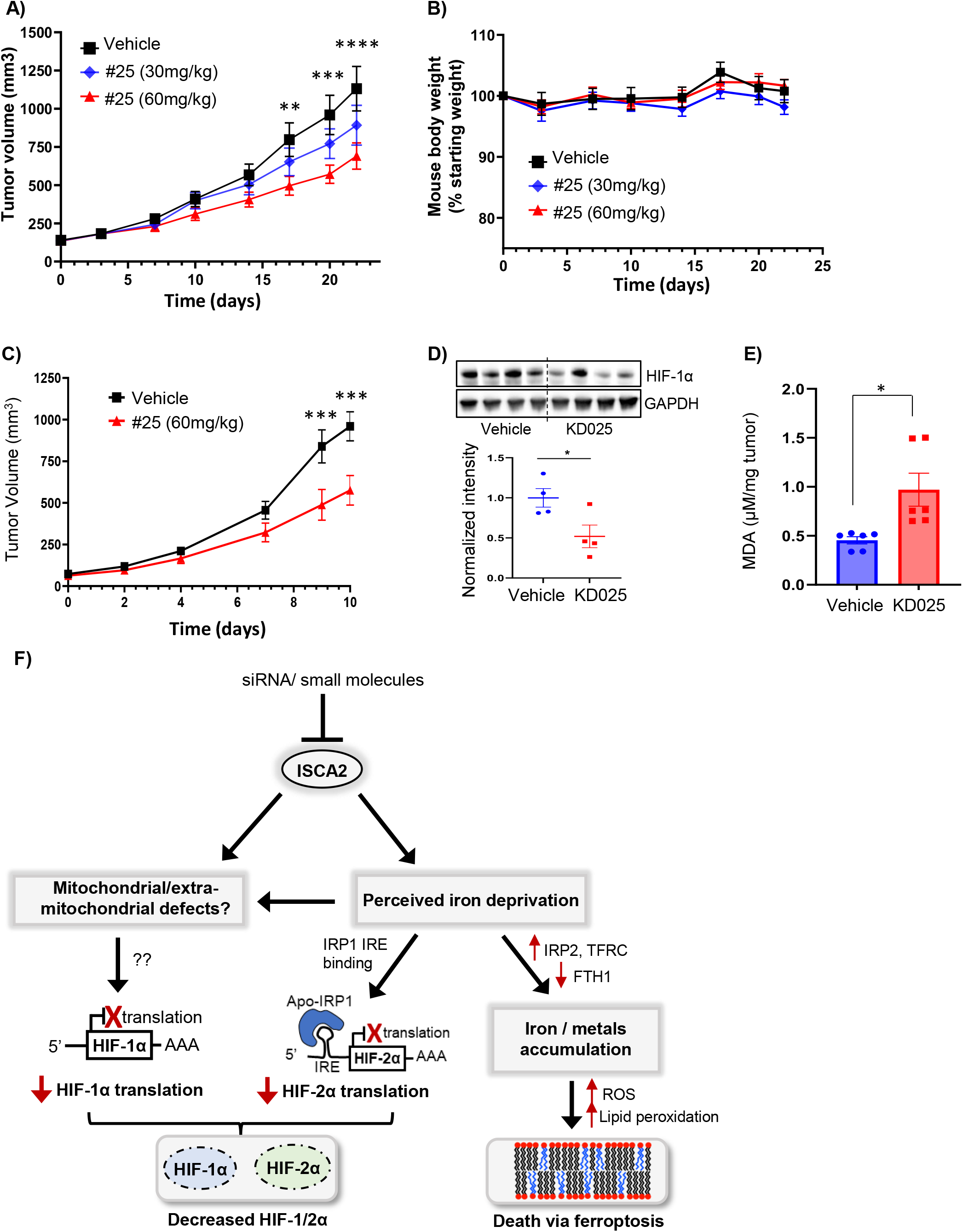
ISCA2 inhibition inhibits xenograft growth *in vivo*. A, B) Effect of treatment of 786-0 subcutaneous xenografts with vehicle or indicated doses of compound #25 (7-8 mice/group) administered orally once daily, on tumor volume (A), or mouse body weights (B). ** indicates p < 0.01; *** p < 0.001; ****p < 0.0001 from Students’ T-tests of vehicle versus 60mg/kg treated group. P NS for 30mg/kg treated group versus vehicle. C) Effect of treatment of RENCA subcutaneous xenografts in syngeneic Balb/c mice with vehicle or 60mg/kg #25 (8 mice/group) once daily. D) Western blots showing HIF-1α and GAPDH with densitometric values normalized to GAPDH below. E) MDA content of RENCA tumors from mice treated with vehicle or 60mg/kg #25. F) Proposed model for effects of ISCA2 blockade. Inhibition of ISCA2 using either small molecules or siRNA results in perceived iron deprivation, which induces upregulation of IRP2, TFRC and downregulation of FTH1, which together promote iron/metals uptake that triggers cell death via ferroptosis. This perceived iron deprivation also drives a shift to the IRE-binding form of IRP1 which inhibits HIF-2α translation. ISCA2 inhibition may also cause other mitochondrial/extramitochondrial defects dependent or independent of perceived iron deprivation that inhibits the translation of HIF-1α through unknown mechanism(s). Thus, ISCA2 inhibition depletes both HIF-1α and HIF-2α and promotes cell death via ferroptosis.

## Discussion

Proteins with Fe-S cofactors play important roles in fundamental cellular processes, including redox reactions, catalysis, protein translation, and DNA synthesis and repair [57]. The biogenesis of Fe/S proteins is catalyzed by complex, conserved assembly systems [43]. In eukaryotes including humans, Fe-S maturation is initiated in mitochondria by the ISC machinery for the maturation of both mitochondrial and extra-mitochondrial Fe-S proteins [58]. ISCA1, ISCA2 and IBA57 play central roles in the later steps of mitochondrial [4Fe-4S] assembly, i.e. in the conversion of [2Fe-2S] clusters to [4Fe-4S] clusters for incorporation into mitochondrial [4Fe-4S] proteins, through a reaction that has not been well defined [44, 59]. Although ISCA1, ISCA2 and IBA57 form a ternary complex to convert two [2Fe-2S] clusters to a [4Fe-4S] cluster in *S. cerevisiae*, this ternary complex is not detected in human cells but instead, biophysical evidence suggest that the ISCA2-IBA57 complex coordinates the transfer of the [2Fe-2S] cluster from the core ISC to the ISCA2-ISCA1 complex for [4Fe-4S] assembly [59-63]. Strikingly, although all three were required for the proliferation of HeLa cells, only ISCA1 is essential for [4Fe-4S] assembly in non-dividing cells [44, 61].

In this study, through efforts to identify selective inhibitors for HIF-2α, we identify ISCA2 as an essential factor for the regulation of both HIF-1α and HIF-2α, and in the maintenance of iron homeostasis in renal cancer cells, and likely in other cancer cell types (**S1A**). We show that ISCA2 inhibition either pharmacologically or using siRNA blocks translation of both HIF-1α and HIF-2α, with greater potency of HIF-2α translational inhibition, by blocking IRE-dependent HIF-2α translation. The mechanism by which ISCA2 inhibition inhibits HIF-1α translation in unclear. However, it is likely that the impact of ISCA2 inhibition at least on HIF-2α translation is due to the triggering of the iron starvation response that promotes the binding of IRP1 to the HIF-2α IRE due to the loss of IRP1’s Fe-S cluster [64]. The iron starvation response is clearly observed with pharmacological inhibition of ISCA2, which results in increased IRP2 and TFRC (increasing iron uptake) and decreased FTH1 (to decrease iron storage; **Fig. 1E-F**). These effects are less apparent with ISCA2 siRNA, possibly due to the increased duration of ISCA2 knockdown required for loss of ISCA2 protein, although shorter duration of ISCA2 knockdown results in marked loss of FTH1 and in HIF-1/2α suggesting that ISCA2 knockdown triggers the iron starvation response prior to when ISCA2 protein is lost (**Fig. 3B**). Significantly, both pharmacological and siRNA-mediated knockdown is associated with significant accumulation of iron and of other transition metals (**Fig. 2E-G, 3H, S2, S3B**). Thus, the decrease in IRP1, IRP2 and TFRC seen after prolonged knockdown of ISCA2 could be a response to this iron/metals accumulation (**Fig. 3A**). Of note, iron/metals accumulation was not observed when other components of the L-ISC, ISCA1 and IBA57 were knocked down, suggesting that ISCA2 possesses unique roles independent of its role in the L-ISC (**Fig. 3H, S3B**). Intriguingly, shRNA knockdown of NFS1, a sulfur donor in ISC biogenesis, also induces IRP2 and TFRC and depletes FTH1, sensitizing cells to ferroptosis (effects on HIF were not investigated) reminiscent of that seen in our current study [65]. Since NFS1 contributes to the maturation of both cytosolic and mitochondrial Fe-S proteins (unlike ISCA1, ISCA2 and IBA57 that are believed to contribute only to mitochondrial [4Fe-4S] assembly), our findings suggest that ISCA2 may play an earlier role in ISC assembly than previously thought [44, 66]. Our findings resemble that of a previous study that identified inhibitors of IRE-dependent HIF-2α translation using a screen in 786-0 cells [67]. Although the molecular target of these inhibitors was not identified (these inhibitors are structurally unrelated to the molecules described in our current study), the investigators also noted similar IRE-independent effects in inhibiting HIF-1α translation [67]. Taken together with our study, these findings suggests that HIF-1α translation may be iron regulated in an IRE-independent manner.

Our data suggest that inhibition of ISCA2 both decreases HIF-α levels and induces ferroptosis by triggering pathways that are independent of ISCA2’s role in mitochondrial [4Fe-4S] assembly (**Fig. 6E**). Although molecular markers that distinguish ferroptosis from other forms of cell death have not yet been identified, we confirmed ferroptosis as the mechanism of cell death by the attenuation of cell death by co-treatment with the iron chelator, DFO, and the ferroptosis inhibitor, liproxstatin but not with the caspase (apoptosis) inhibitor, ZVAD-FMK (**Fig. 4G-H**) [68]. Furthermore, cells transfected with ISCA2 siRNA showed increased numbers of dead cells with increased cell size and plasma membrane rupture commonly observed in ferroptotic cells and showed changes in cell membrane permeability associated with both apoptosis and necrosis that resembled that or RSL3 (**S5, Fig. 4E-F**) [49]. ISCA2 levels are decreased with pVHL loss and patients with low ISCA2 levels have significantly decreased overall survival (**Fig. 5C, H, I**) suggesting that ISCA2 may play a tumor suppressor role. Paradoxically, pVHL deficient cells with lower levels of ISCA2 are more sensitive to death induced by ISCA2 inhibition (**Fig. 5D-E**), which could be rescued by ISCA2 overexpression (**Fig. 5F**). Taken together, the data suggest a therapeutic index for inducing cell death through ISCA2 inhibition specifically in pVHL deficient cells. Significantly, we show that treatment with #25 by oral administration, significantly inhibits the growth of both 786-0 and RENCA tumor xenografts, decreases HIF-1α levels, increases lipid peroxidation, and is well tolerated at the therapeutic dose (**Fig. 6A-E**), providing proof-of-concept for the efficacy and safety of ISCA2 inhibition for the treatment of ccRCC and other HIF driven and/or ferroptosis susceptible cancers.

## Materials and methods

### High Throughput Screen

The primary HTS was performed using 786-0 cells stably expressing HRE-Luciferase containing 5X copies of the HRE from the VEGFA in the pGL3 vector backbone. These cells were used to screen the entire MLSMR of ∼360,000 compounds at 18.8 μM in 1536-well format. The assay performed robustly with a Z’ average over 272 plates of 0.84, and a signal-to-background ratio of 76.8. Summary of the methods and results of the screen and counterscreens are described in Pubchem AIDs 624357, 624352, 652580, 652581 and 651589. Compound #1 was purchased from Molport (Beacon, NY) CAS ID: 862974-22-9, whereas #25 was synthesized as described (in press).

### Cell Lines and transfection reagents

ACHN, MIAPaCa-2 and 786-0 cells were from ATCC (Manassas, VA), whereas RCC4 cells were from M. Celeste Simon (University of Pennsylvania). All commercially available cell lines were authenticated using STR fingerprinting upon receipt and stored in frozen aliquots. Fresh vials were thawed for use in experiments and discarded after 30-35 passages or approximately 2 months. Mycoplasma testing using the MycoAlert detection kit (Lonza, Ben OR) was performed every 2 months. Cells were maintained at 37°C 5% CO_2_ in Dulbecco’s MEM (Life Technologies, ThermoFisher Scientific, Waltham MA) with 10% FBS. Hypoxia incubations were performed using a Whitley H35 Hypoxystation (HypOxygen, Frederick, MO). Lipofectamine™ RNAiMAX Transfection Reagent (ThermoFisher) was used for siRNA transfection. siRNAs were purchased from Dharmacom (Horizon Discovery, Lafayette CO): siCon (D-001810-10) siISCA1 (L-014678-02), siISCA2#1 (L-019329-01), siISCA2#2 (L-019329-02), siIBA57 (L-021938-01).

### Animal studies

NRG or Balb/c mice (10 per group) were injected subcutaneously with 10^7^ 786-0 cells or 2 × 10^6^ RENCA cells respectively. When tumors became established, mice were stratified into groups of approximately equal tumor sizes (average 150mm^3^ for 786-0 and 100mm^3^ for RENCA) and dosing was initiated. Mice were dosed via PO daily with vehicle (0.5% methyl cellulose, 1% Tween 80) or indicated doses of #25 (HCL salt). Tumor diameters were measured twice weekly at right angles (d_short_ and d_long_) using electronic calipers and tumor volumes calculated by the formula volume = (d_short_)^2^ x (d_long_)^2^ [69]. Studies were terminated when tumors reached 1000mm^3^. All animal were conducted in compliance with relevant local guidelines and were approved in the Institutional Animal Care and Use Committee.

### TaqMan Quantitative RT-PCR

Cell lysis and RNA extraction from siRNA transfected cells were performed using the RNeasy RNA extraction kit (Qiagen Inc, Germantown, MD). 1 µg of total RNA was reverse transcribed with the High-capacity RNA-to-cDNA kit (Applied Biosystems, Foster City, CA). For RT-PCR, cDNA was combined with TaqMan Gene Expression Mastermix (Applied Biosystems), and TaqMan primers for *B2M* (ThermoFisher, Hs00187842_m1), *HIF1A* (Hs00153153_m1), *EPAS1* (Hs01026149_m1), POU5F1 (Hs03005111_g1) or *SOX2* (<u>Hs04234836_s1</u>). RT-PCR experiments were run on a QuantStudio 6 Flex System (Applied Biosystems). mRNA levels were calculated relative to the housekeeping gene, B2M using the ΔΔCt method as recommended by the manufacturer.

### Immunoblotting

Western blotting was performed as previously described [70-72]. Antibodies for western blotting are listed in **Supplemental methods**. Images were acquired using Fluorochem M imaging system (Protein Simple, San Jose, CA). Blot quantification was performed using multiplex band analysis provided by the AlphaView Software (Protein Simple).

### Human subjects and ISCA2 IHC

Studies were conducted in accordance with the Declaration of Helsinki. Human ccRCC and uninvolved kidney tissue were obtained from archival samples from patients who had provided written informed consent according to protocols approval by the institutional review boards at HCI. TMAs constructed from formalin fixed paraffin embedded tumor tissue from advanced metastatic RCC treated with antiangiogenic therapy. Overall survival was defined as time from initiation of therapy until death. All specimens were reviewed by pathologists and representative tumor areas identified. Three spatially separated 2mm cores from archival FFPE tissue were included for each case. TMAs and tissue were stained for ISCA2 (HPA030492 Atlas antibodies, Bromma, Sweden) using conditions optimized using normal kidney according to the manufacturer’s protocols using the BenchMark Ultra automated slide stainer (Ventana Medical Systems, Roche, Oro Valley, AZ).

### Statistical analysis

P-values, curve fitting and IC_50_ calculations, Kaplan Meier plots (with logrank p-values and hazard ratios) were determined using Prism Version 9.3.1 (GraphPad Software, San Diego). Where appropriate differences between groups was determined using Students’ T-tests using p < 0.05 as statistically significant.

Details on the DARTS, thermal shift and other assays as provided in **Supplemental Data**.

## Acknowledgements and funding sources

Supported by NIH grants R03DA033980, CA181106 and CA217905 and Dod KCRP grant KC190022 to MYK and NIH Hematology Training Program Grant T32DK007115 to DF. We acknowledge financial support by the Huntsman Cancer Foundation. Research reported in this publication utilized the Preclinical Research Shared Resource at Huntsman Cancer Institute at the University of Utah. The content is solely the responsibility of the authors and does not necessarily represent the official views of the NIH.

