## Supplemental Figures for "ISCA2 inhibition decreases HIF and induces ferroptosis in clear cell renal carcinoma"

**Supplemental Figure 1**

A)

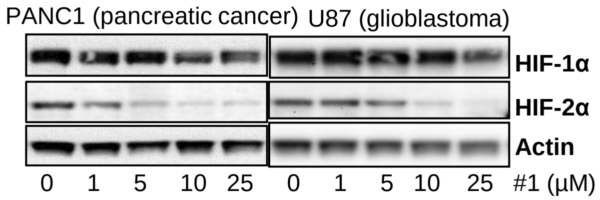


B)

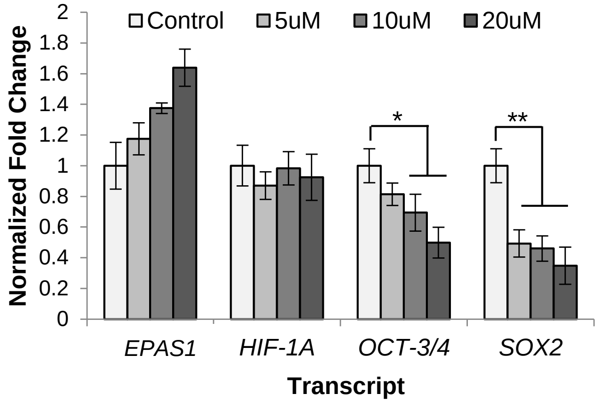


C)

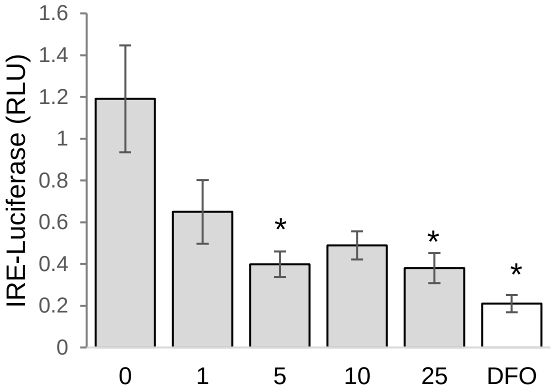


D)

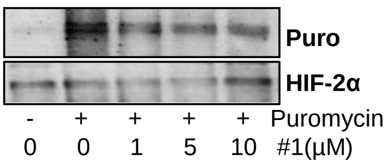


Supplemental Fig 1: A) Western blots showing the effects of 24 hours' treatment of PANC1 and U87 cells with indicated concentrations of compound #1. Cells were exposed to hypoxia (1%$O_2$) for 24 hours to induce HIF expression. B) Quantitative RT-PCR showing effects of #1 on the transcription of *HIF1A, HIF2A* (*EPAS1*) and HIF-2 target genes *OCT-3/4* (*POU5F1*) and *SOX2* in hypoxic ACHN cells. Effect of compound #1 on luciferase activity driven by a HIF-2α Iron-Responsive Element (IRE)-luciferase reporter construct transiently transfected into ACHN cells. * $p < 0.05$. D) Western blot showing puromycin incorporation into immunoprecipitated HIF-2α in hypoxic ACHN cells treated with #1 for 24 hours.

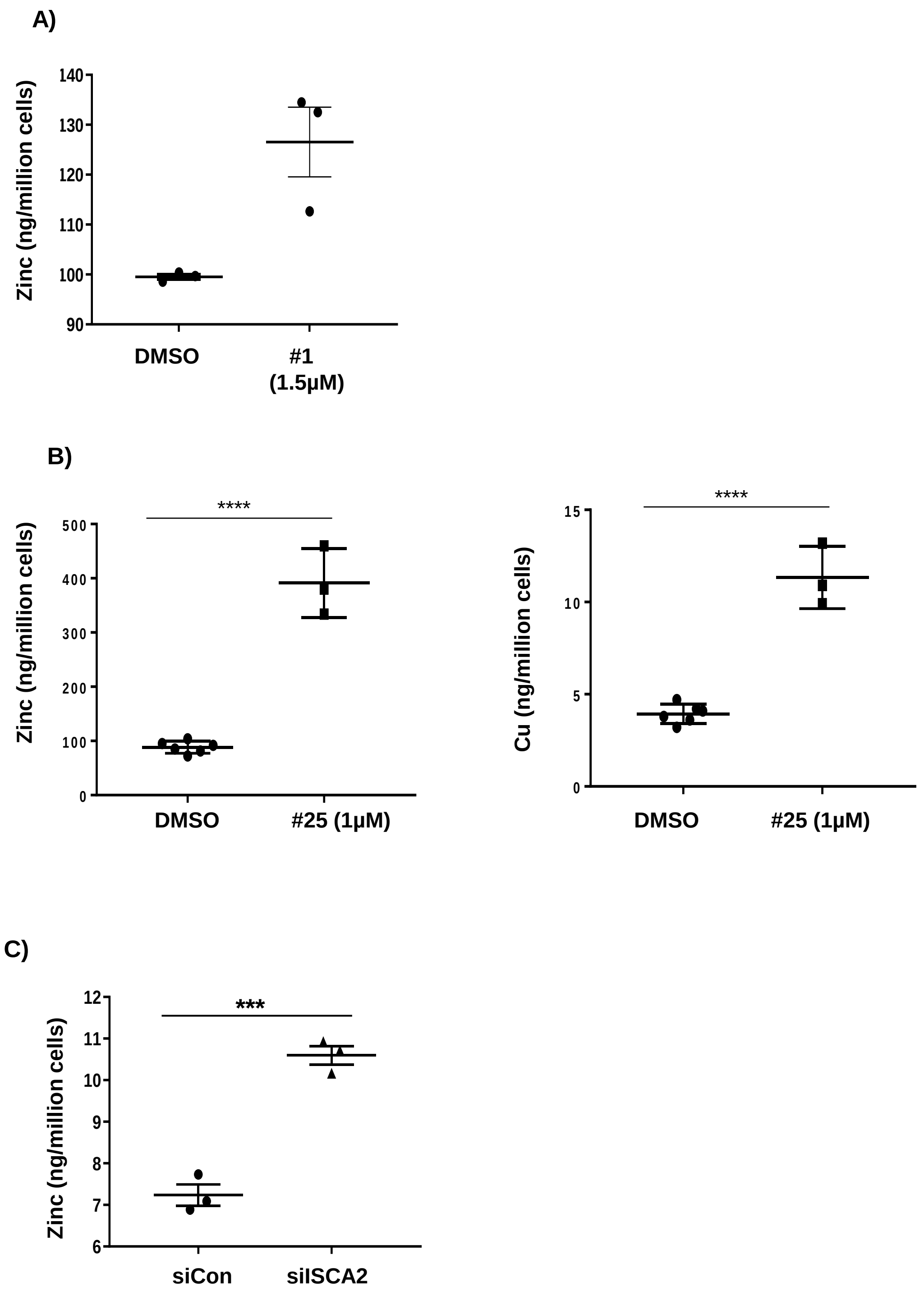


Supplemental Fig 2: Metals content of 786-0 cells treated with; A) #1 (24 hours), B) #25 (24 hours) or C) Control or ISCA2 siRNA (6 days) as determined using ICP-MS.

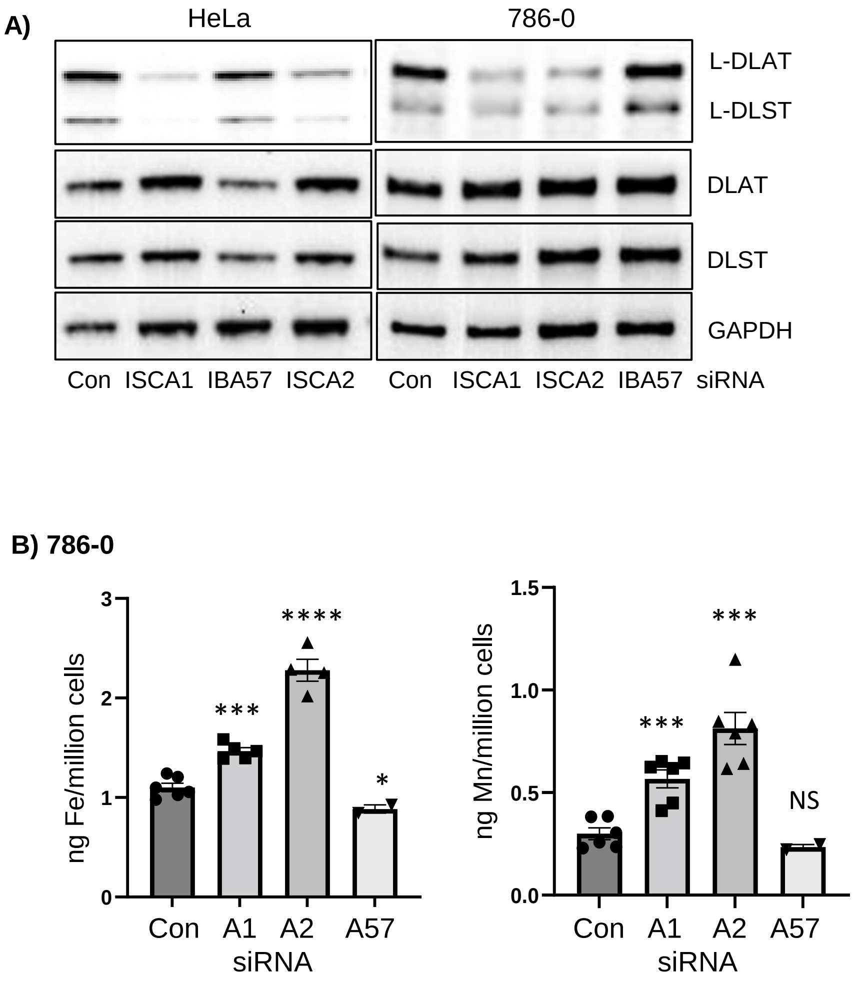


Supplemental Fig 3: A) Western blots showing effect of indicated siRNA transfections (6 days) on lipoylation of DLAT and DLST in HeLa and 786-0 cells. B) Effect of 5-day siRNA knockdowns on metals accumulation in 786-0 cells detected using ICP-MS. Approximately equal cell numbers were submitted for analysis. All data are representative or averages of at least two independent experiments ±SEM.

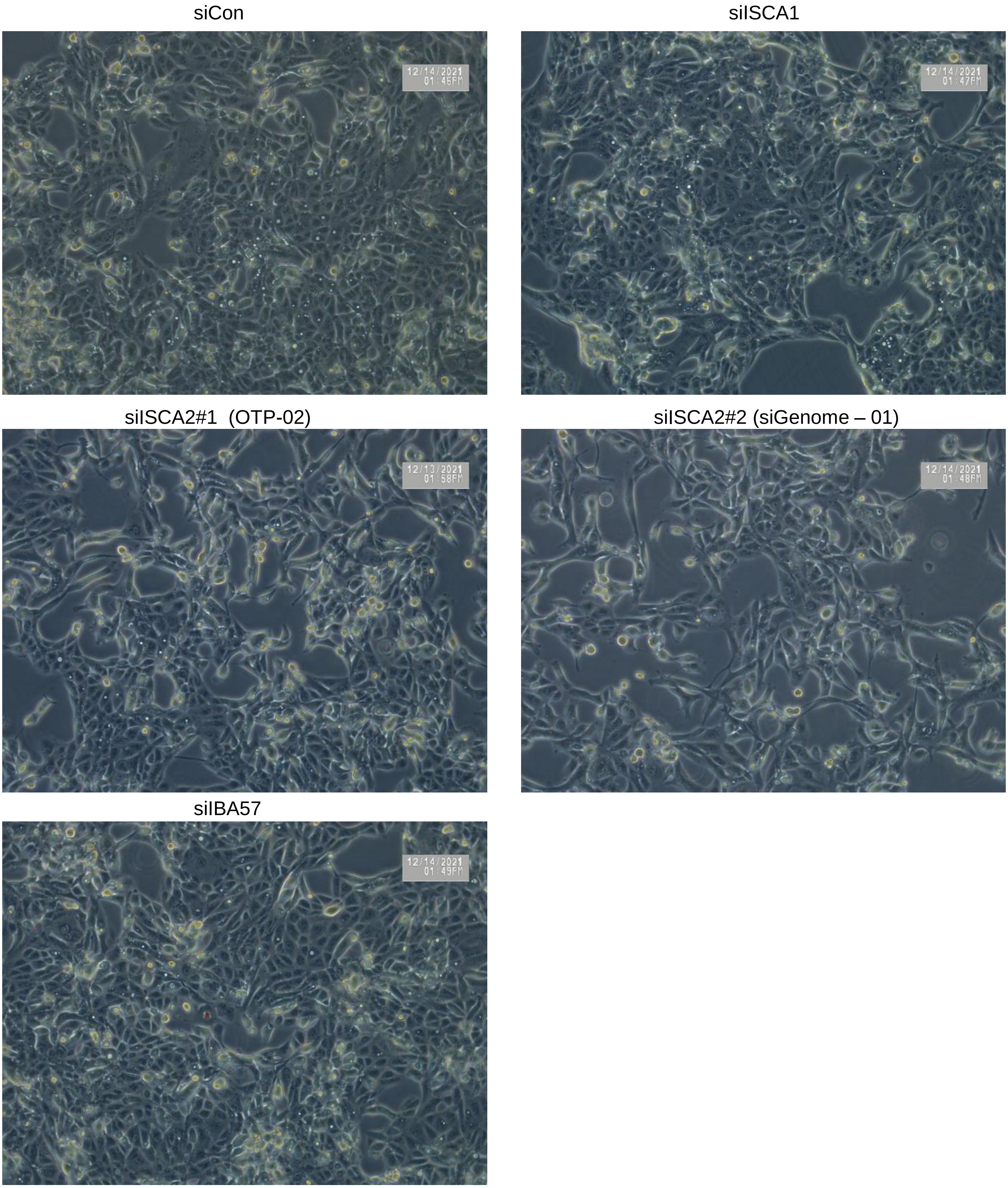


Supplemental Fig 4: Representative phase contrast photomicrograph of adherent ACHN cells in flasks after 5 days transfection with indicated siRNAs used for ICP-MS analysis.

**Supplemental Figure 5**

siCon

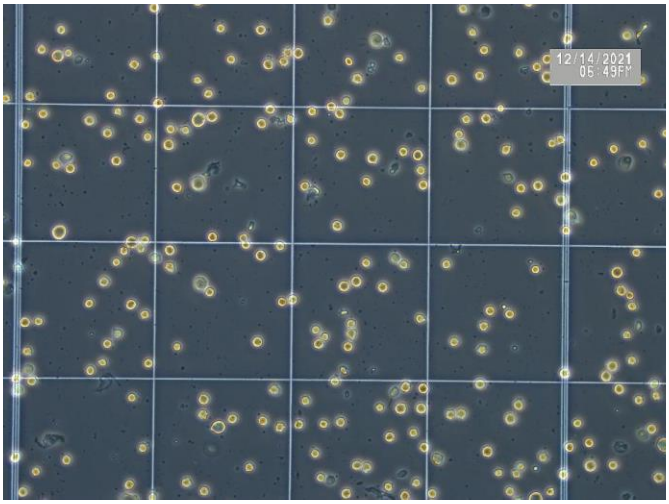

siISCA1

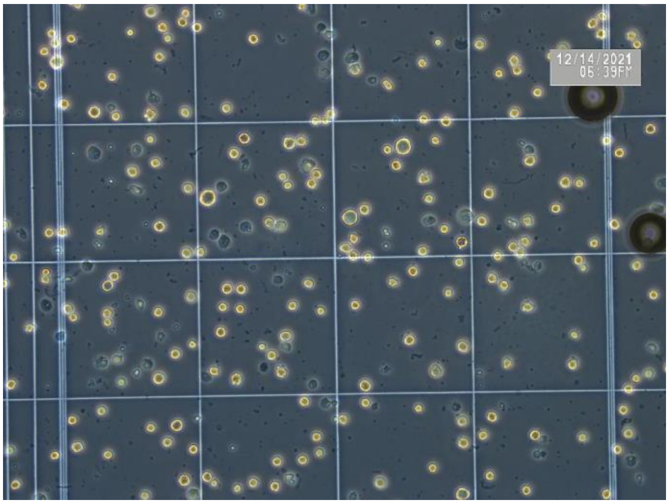

sISCA2

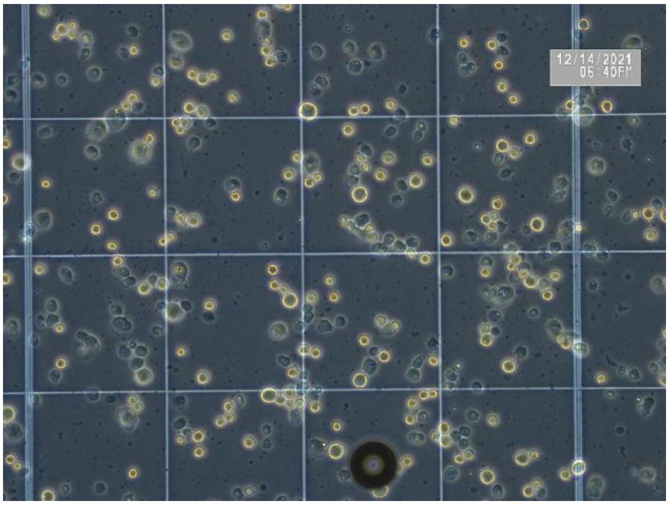

siIBA57

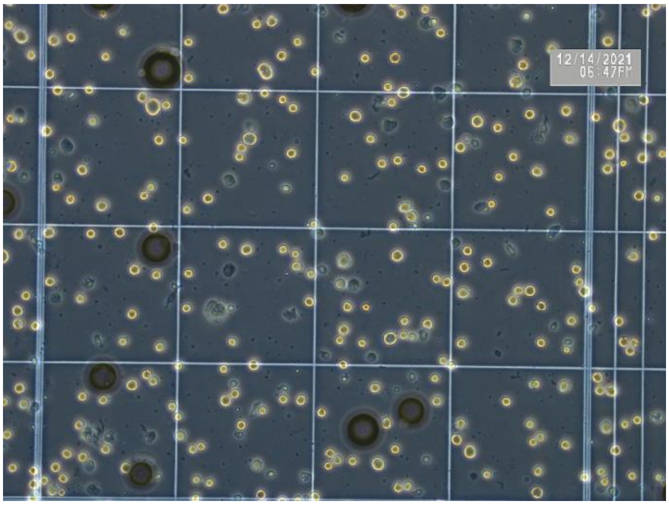

Supplemental Fig 4: Representative phase contrast photomicrographs within hemocytometer counting chambers of trypsinized ACHN cells after 5 days transfection with indicated siRNAs used for ICP-MS analysis. Images indicate the increased size and reduced number of cells transfected with siISCA2

**Supplemental Figure 6**

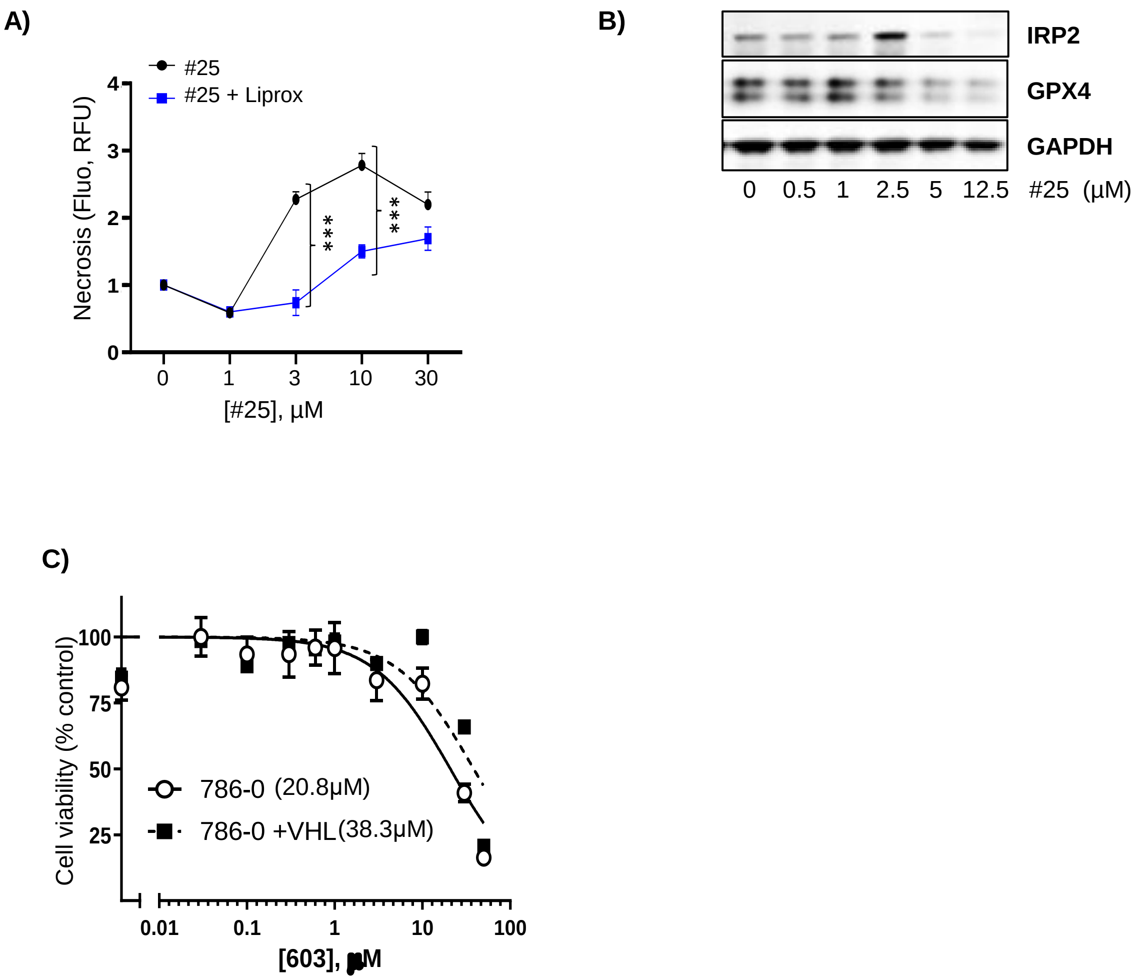


Supplemental Fig 6: A) Dose response effect of #25 ± liproxstatin (1µM) on inducing loss of membrane permeability (necrosis) after 72hrs' treatment. B) Western blot showing effects of treatment with #25 on GPX4 and IRP2 at higher concentrations. Blots show reductions in IRP2 and GPX4 at ≥5µM treatment. C) Resazurin cell viability assay of 786-0 parental or 786-0 cells with pVHL reconstitution treated with #1 for 72 hours. Average $IC_{50}$ values (µM) are shown in brackets.
