## Supplemental methods for "ISCA2 inhibition decreases HIF and induces ferroptosis in clear cell renal carcinoma"

### Antibodies for western blotting

Membranes were probed with primary antibodies for HIF-1α (BD Biosciences, 610959, Franklin Lakes, NJ), HIF-2α (Cell Signaling Technologies (CST), 70862, Danvers MA), ISCA1 (Abnova, H00081689-M01, Walnut, CA ), ISCA2 (Genetex, GTX45528, Irvine, CA or Bethyl Laboratories, A305-843A-M, Montgomery, TX), IBA57 (Abcam, ab180161, Waltham, MA), Ferritin heavy chain (Santa Cruz Biotechnology (SCB), sc-376594, Dallas, Texas), IRP1 (SCB, sc-14216), IRP2 (SCB, sc-33682), Transferrin Receptor (CST, 13113), Actin (CST, 3700), GAPDH (CST, 5175), Puromycin (MilliporeSigma, MABE343, St. Louis, MO), COX IV (Abcam, ab16056), DLAT (CST, 12362), and DLST (CST, 11954)

### DARTS assay

*Sample preparation.* Prior to digestion, protein concentration was determined using bicinchoninic acid (BCA) protein assay (Pierce, Waltham, MA). Samples were then digested using a modified Filter-aided Sample Preparation (FASP) protocol (1). In brief, each sample was transferred to a 3-kDa molecular weight cutoff filter (Millipore, MA, USA). All following centrifugations were carried out at 14,000 rpm in a benchtop microfuge. First, the samples were washed 3x with UA buffer (8M urea, 50 mM ammonium bicarbonate), and cysteine disulfide bonds were reduced with 5 mM dithiothreitol (DTT) at 30°C for 60 min followed by cysteine alkylation with 15 mM iodoacetamide (IAA) in the dark at room temperature for 30 min. Following alkylation, the samples were washed 2x with UB buffer (1 M urea, 50 mM ammonium bicarbonate). The samples were finally transferred to a tube and subjected to overnight digestion with mass spec grade Trypsin/Lys-C mix (Promega, Madison, WI). Digested protein samples were finally desalted using a $C_{18}$ TopTip (PolyLC, Columbia, MD) according to the manufacturer's recommendation), and the organic solvent was removed in a SpeedVac concentrator prior to LC-MS/MS analysis.

*LC-MS/MS analysis.* Dried samples were reconstituted in 100mM ammonium formate pH ~10 and analyzed by 2DLC-MS/MS using a 2D nanoACQUITY Ultra Performance Liquid Chromatography (UPLC) system (Waters corp., Milford, MA) coupled to an Orbitrap Elite mass spectrometer (Thermo Fisher Scientific). Peptides were loaded onto the first dimension column, XBridge BEH130 $C_{18}$ NanoEase (300 µm x 50 mm, 5 µm) equilibrated with solvent A (20mM ammonium formate pH 10, first dimension pump) at 2 µL/min. The first fraction was eluted from the first dimension column at 12% of solvent B (100% acetonitrile) for 4 min and transferred to the second dimension Symmetry $C_{18}$ trap column 0.180 x 20 mm (Waters corp., Milford, MA) using a 1:10 dilution with 99.9% second dimensional pump solvent A (0.1% formic acid in water) at 20 µL/min. Peptides were then eluted from the trap column and resolved on the analytical $C_{18}$ BEH130 PicoChip column 0.075 x 100 mm, 1.7µm, particles (NewObjective, MA) in a 96-min gradient of 2-28% solvent B at a flow rate of 400nL/min. The following 9 first dimension fractions were eluted at 13, 14, 15, 16, 17.5, 19, 21, 23, and 60% solvent B. The mass spectrometer was operated in positive data-dependent acquisition mode. MS1 spectra were measured with a resolution of 60,000, an AGC target of 1e6 and a mass range from 350 to 1400 m/z. Up to 10 MS2 spectra per duty cycle were triggered, fragmented by collisium-induced dissociation, and acquired in the ion trap with an AGC target of 1e4, an isolation window of 2.0 m/z and a normalized collision energy of 35. Dynamic exclusion was enabled with duration of 20 sec.

*Data analysis.* All mass spectra from were analyzed with MaxQuant software version 1.5.5.1. MS/MS spectra were searched against the *Homo sapiens* Uniprot protein sequence database (version July 2015) and GPM cRAP sequences (commonly known protein contaminants). Precursor mass tolerance was set to 20ppm and 4.5ppm for the first search where initial mass recalibration was completed and for the main search, respectively. Product ions were searched with a mass tolerance 0.5 Da. The maximum precursor ion charge state used for searching was 7. Carbamidomethylation of cysteines was searched as a fixed modification, while oxidation of methionines and acetylation of protein N-terminal were searched as variable modifications. Enzyme was set to trypsin in a specific mode and a maximum of two missed cleavages was allowed for searching. The target-decoy-based false discovery rate (FDR) filter for spectrum and protein identification was set to 1%.

### Luciferase assays

786-0 and MIAPaCa-2 cells stably expressing HRE-luciferase reporter and 786-0 cells stably expressing the HIF-2α IRE-luciferase reporter, were seeded at appropriate densities for transfection or treatment in 96-well plates and were transfected with siRNA or treated with drugs for the indicated time period. Luciferase assays were performed using the Steady-Glo Luciferase Assay System (Promega, E2510) in 96-well plates according to the manufacturer's protocol and luminescence was measured using a plate reader (Victor™ X3, PerkinElmer, Waltham, MA).

**Cell viability assays**

For effects of treatment with small molecules, 786-0 cells were seeded at 4,000 cells/well in 96-well plates and allowed to adhere overnight after which cells were treated with a concentration range of small molecules. Cell viability was determined after 24hours' treatment using the resazurin (AR002, R&D Systems, Minneapolis, MN). For siRNA transfections, 786-0 cells were seeded in 96-well plates at 1,500 cells per well, and siRNAs were transfected in the next day. Apoptosis and necrosis detection reagents from the RealTime-Glo™ Annexin V Apoptosis and Necrosis Assay kit (Promega, JA1011) were added 48 hours after siRNA transfection, and recombinant human TRAIL (Biolegend, San Diego, CA) was added as a positive control of apoptosis and necrosis, according to the manufacturer's protocol. Luminescence and fluorescence were measured using a plate reader at several time points throughout the study up to 96 hours' after siRNA transfection.

**Puromycin labelling and Immunoprecipitation (IP)**

Newly synthesized proteins were labelled with puromycin using the SUnSET assay as described previously (2). Briefly, cells were incubated with 1µg/mL puromycin (Gibco, A1113803) for 30 minutes and then incubated for an additional hour with fresh media at 37°C in 5% $CO_2$ incubator. IP was performed using the HIF-1α (BD Biosciences, 610959) and HIF-2α (Novus Biologicals, NB100-122) antibodies as described previously (3). Briefly, 1 mg lysate in lysis buffer was precleared with protein A-Sepharose beads (Sigma-Aldrich, P3391) for 1 hour at 4°C. The precleared lysate was incubated with an antibody for 2~3 hours at 4°C and then further incubated after adding protein A-Sepharose beads overnight at 4°C. The beads were washed with lysis buffer and then the immunoprecipitated proteins were analyzed using immunoblotting.

**Inductively coupled plasma mass spectrometry (ICP-MS)**. 786-0 or ACHN cells were seeded at appropriate densities for transfection or treatment in T 75cm$^2$ flasks to obtain ≥ 2 million cells per replicate with each experiment performed in triplicate. Cells allowed to adhere overnight, then treated with compounds in DMSO or DMSO alone (vehicle) for a further 24 hours, or transfected with siRNA for 5 days, after which cells were detached by trypsinization, counted, washed twice in PBS and pelleted. Experiments were performed using three replicate T75cm2 flasks per condition. A 5:1 mixture of nitric acid (OPTIMA Grade, 70%, Fisher Scientific) and ultrapure hydrogen peroxide (ULTREX II, 30%, Fisher Scientific) was added to cell pellets. This mixture was allowed to digest overnight, heated until dry, and resuspended in 2% nitric acid for analysis using an Agilent 7900 ICP-MS (Agilent Technologies, Santa Clara, CA). Calibration standard solutions for determination of Fe were prepared from Agilent multi-element calibration standard-2A. An Agilent Environmental Calibration Standard was used as an independent control. PBS-only control digestions were used to measure background. Metal readings were normalized to cell number.

**Thermal Shift assays**

Thermal shift assays were performed by monitoring the change in protein melting temperature (Tm) in the absence or presence of test compounds, using the hydrophobic protein binding dye, SYPRO Orange (S6650, Thermo Fisher Scientific, Waltham, MA), measured using the LightCycler 480 (Roche Life Sciences, Indianapolis, IN) according to the manufacturer's protocol. Recombinant ISCA2 was produced by expressing amino acid residues 9-154 of ISCA2 (ISCA2 lacking its mitochondrial localization sequence) in the pET28 vector containing an N-terminal His6 tag in Rosetta (DE3) competent cells (Novagen, Millipore Sigma). ISCA2 production was induced by treating ISCA2 transformed log phase cells with 0.25mM IPTG for 4 hours at 18oC. ISCA2 was purified using Ni2+ affinity purification according to standard protocols and eluted in 50mM Tris-Cl pH7.4, 150mM NaCl, 5mM DTT. Thermal shift assays were performed in 384-well plates using 1µl of a 10X concentration of SYPRO Orange, 8µl of ISCA2 (4µg protein) and 1µl of 1-2mM stock of test compound per well. The LightCycler was used according to the following setup: LightCycler 480 Instrument Temperature Setup: First target of 20°C, with a Hold of 15 seconds; second target of 95°C, with Acquisition Mode of Continuous, and 10 acquisitions per degree C; and third target of 20°C, with a Hold of 15 seconds. Melting temperature $T_m$s were determined using Roche Protein Melting Analysis Software.

**ISCA2 IHC**

Briefly, heat-induced antigen retrieval was performed in pH 8.5 buffer for 36 minutes at 95C, and the ISCA2 antibody was applied at 1:200 dilution for 80 minutes at 37C. An amplification kit was applied to increase signal and detection was performed using a DAB kit with a 12-minute counterstain with hematoxylin. ISCA2 staining intensity was quantitated using the Halo Digital Imaging Platform (Indica Labs, Albuquerque, NM). KM plots were plotted using GraphPad Prism software 9.3.1 (GraphPad Software, San Diego CA).

**Lipid peroxidation assays**

Malondialdehyde (MDA), an indicator of lipid peroxidation, was quantitated in cells and tumor tissue using the Thiobarbituric Acid Reactive Substances (TCA Method) (Cat 700870, Cayman Chemicals, Ann Arbour Michigan) according to the manufacturer's protocol. C11-BODIPY (581/591) was purchased from ThermoFisher and used to assess lipid peroxidation in #25-treated cells according to the manufacturer's protocol.
